# A fast blind source separation algorithm for decomposing ultrafast ultrasound images into spatiotemporal muscle unit kinematics

**DOI:** 10.1101/2022.11.22.517488

**Authors:** Robin Rohlén, Jonathan Lundsberg, Nebojsa Malesevic, Christian Antfolk

## Abstract

**Objective:** Ultrasound can detect individual motor unit (MU) activity during voluntary isometric contractions based on their subtle axial displacements. The detection pipeline, currently performed offline, is based on displacement velocity images and identifying the subtle axial displacements. This identification can preferably be made through a blind source separation (BSS) algorithm with the feasibility of translating the pipeline from offline to *online*. However, the question remains how to reduce the computational time for the BSS algorithm, which includes demixing tissue velocities from many different sources, e.g., the active MU displacements, arterial pulsations, bones, connective tissue, and noise.

**Approach:** This study proposes a fast velocity-based BSS (velBSS) algorithm suitable for online purposes that decomposes velocity images from low-force voluntary isometric contractions into spatiotemporal components associated with single MU activities. The proposed algorithm will be compared against stICA, i.e., the method used in previous papers, for various subjects, ultrasound- and EMG systems, where the latter acts as MU reference recordings.

**Main results:** We found that the spatial and temporal correlation between the MU-associated components from velBSS and stICA was high (0.86 ± 0.05 and 0.87 ± 0.06). The spike-triggered averaged twitch responses (using the MU spike trains from EMG) had an extremely high correlation (0.99 ± 0.01). In addition, the computational time for velBSS was at least 50 times less than for stICA.

**Significance:** The present algorithm (velBSS) outperforms the currently available method (stICA). It provides a promising translation towards an online pipeline and will be important in the continued development of this research field of functional neuromuscular imaging.

## Introduction

Recording motor unit (MU) activity in humans is based on electromyography (EMG) techniques such as needle EMG [1–3] and surface EMG [4,5]. These techniques record *temporal electrophysiological* MU activity, e.g., the action potential and the neural discharges. Although the scanning-EMG has previously been used to study *spatial* distributions of a MU in a 1D corridor of muscle units [6], these techniques have very limited information for a large field of view in the muscle. In addition, these techniques cannot obtain *mechanical* activity induced by the motoneuron. Obtaining the *spatiotemporal mechanical* activity from many MUs simultaneously over a range of force levels enables studying recruitment in terms of spatial positions of individual MUs [7] and their different types (slow, fast resistant, and fast fatigable) [8].

Ultrasound imaging can detect individual MU activity during *voluntary* isometric contractions [7,9–16]. The principle of detecting MU activity is based on the depolarization of the MU fibres, resulting in subtle axial displacements. These subtle displacements of a MU can be detected using three methodological steps performed *offline*. First, plane wave imaging is used to increase the frame rate (> 1 kHz) [17]. Second, displacement velocity images are calculated based on the raw ultrasound (radio frequency) data [18]. Third, the activity of individual MUs is localized in the displacement velocity images through spike-triggered averaging (using neural discharges from EMG concurrently) [16] or using blind source separation (BSS) [7,9–15]. The BSS algorithm extracts spatiotemporal components, where a subset is MUs [14,19].

The advantage of using BSS is that only a few seconds of data can be used for MU identification [11,19]. In contrast, spike-triggered averaging needs long sequences to provide estimates with a high signal-to-noise ratio. Given this, BSS provides the feasibility of translating the methodological pipeline for MU identification from an *offline* to an *online* method. The significant reduction in computational time for the first two steps of the pipeline, i.e., the plane wave imaging, beamforming, and velocity calculations, can be performed using graphics processing unit (GPU) programming like the setup in ultrasound location microscopy of the brain and kidney [20,21]. However, the question remains how to reduce the computational time for the third step of the pipeline, the BSS algorithm. The input to the BSS algorithm is 2D images over time with mixtures of tissue velocities from many different sources, e.g., the active MU displacements, arterial pulsations, bones, connective tissue, and noise. The challenges in implementing a fast BSS method are that the input needs to be whitened [22], which usually means calculating the inverse of a large covariance matrix before estimating a projection vector that separates the 2D images into MU activities and other non-MU components.

This study proposes a fast BSS algorithm suitable for online purposes that decomposes velocity images from low-force voluntary isometric contractions into spatiotemporal components associated with single MU activities. The proposed algorithm will be compared against the method used in previous papers for different research systems, subjects, and EMG systems (as reference). Six ultrafast ultrasound recordings (based on two datasets) from six different subjects will be used. Three datasets have needle EMG as a reference from one ultrasound research system, and the other three have high-density surface EMG as a reference from another ultrasound research system.

## Methods

### Experimental data

The experimental data consists of two datasets with three recordings of the biceps brachii of healthy subjects in each dataset (Fig. 1A). These datasets come from two previous studies using different ultrasound research systems [14,19]. The MU reference is based on two different EMG recordings, i.e., concentric needle EMG and high-density surface EMG where we only select one MU for each recording. Both datasets consist of ultrafast ultrasound imaging to record images from the cross-section of the biceps brachii under low force levels (Fig. 1B). First, displacement velocity images are calculated based on the raw ultrasound data (Fig. 1C). Then, the velocity images are decomposed into components using spatiotemporal independent components analysis (stICA) [14,19] and the proposed velocity-based blind source separation (velBSS) algorithm (Fig. 1D). Finally, out the decomposed components, we select one component for each recording and decomposition algorithm for comparison.

**Figure 1.**
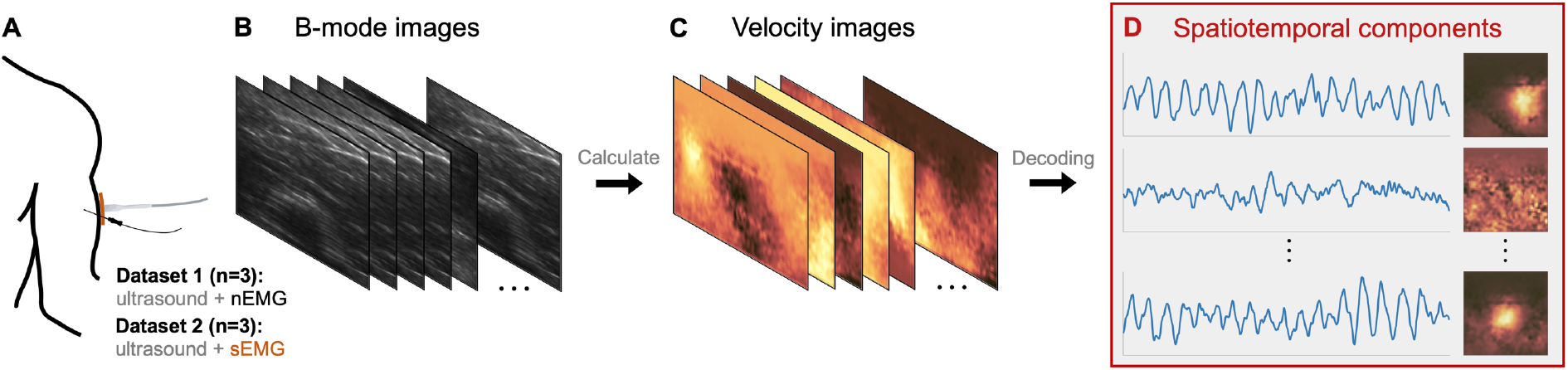
**A.** The experimental data consists of two datasets with three recordings of the biceps brachii of healthy subjects in each dataset. The MU reference is based on two different EMG recordings, i.e., needle electromyography (nEMG) and surface EMG (sEMG). **B.** B-mode images of the cross-section of the biceps brachii under low force levels. **C.** Displacement velocity images are calculated based on the raw ultrasound data. **D.** The velocity images are decomposed into components using spatiotemporal components analysis (stICA) and the proposed velocity-based blind source separation (velBSS) algorithm.

#### First dataset with three recordings

Two-second-long recordings of three healthy subjects (28.3 ± 0.6 years old; one male and two females) were retrospectively included from a previous study [19]. A subject was seated on a chair to perform a weak isometric elbow flexion with the elbow flexed at 90°. Before supinating their lower arm, a physician inserted a concentric needle electrode distal to the muscle belly. The subject received feedback through a loudspeaker of the EMG activity and the quality inspection of the physician. When the physician was satisfied with the quality of the obtained EMG signal from one or two MUs, the subject was instructed to hold that position and force level. The ultrasound probe was positioned proximal to needle insertion such that the image plane captured the needle tip (verified by mechanical poking of the needle). The ultrasound and nEMG recordings were synchronised based on a customised procedure using a trigger at the start of the recording such that the data could later be aligned in a post-processing stage [19].

The EMG recordings were performed on a Cadwell Sierra Wave EMG machine (Cadwell Laboratories Inc., Kennewick, WA, USA) sampling at 64 kHz with a 38×0.45 mm concentric needle electrode (AMBU Neuroline, DEN). The nEMG signals were used to identify one MU spike train using template matching [19,23]. The MU action potentials were manually double-checked, and a few superpositions were solved using manual editing.

The ultrasound probe (Ultrasonix L14-5/38 with a centre frequency of 9 MHz) was placed on the muscle belly proximal to the needle insertion to the skin. The radio frequency data was recorded using SonixTouch (Ultrasonix Medical Corporation, Richmond, CA), sampled at 40 MHz with a frame rate of 2 kHz due to single plane wave imaging. Each dataset comprised 4031 frames (2080×128 pixels, each with a field of view equal to 40×40 mm).

After delay-and-sum beamforming, each beam axis (line) was band-pass filtered (2-15 MHz with order 6) [19]. Next, each pixel was filtered over time with a 1D median filter with an order equal to 10 ms (12 samples) and a high-pass filter of 5 Hz [19]. Next, displacement velocity images were calculated based on the radio frequency data using 2D autocorrelation velocity tracking [18] with 1 mm (40 samples) in-depth and a sliding window of 10 ms (20 frames) [19]. Next, the regions were downsampled axially by a factor of sixteen, resulting in 4012 frames where each frame was 128×128 pixels (about 0.3×0.3 mm per pixel). Finally, an ROI of 64×64 pixels was selected [19] for further analysis, where the ROI was placed to contain the activity of a MU.

The study was conducted following the Declaration of Helsinki and the procedure approved by the Swedish Ethical Review Authority (2019-01843). Informed consent was obtained from all participants after receiving a detailed explanation of the study procedures.

#### Second dataset with an additional three recordings

Eight-second-long recordings of three healthy subjects (29.3 ± 4.6 years old; three males) were retrospectively included from a previous study [14]. A subject was seated on a chair with the right arm fixed for elbow joint torque measurement (elbow flexed at 135°). In the three included datasets, each subject was requested to perform a 60-second-long isometric elbow flexion at 5%, 2%, and 2% of the maximum voluntary contraction, which was performed prior to the recording. The sEMG signals were acquired during the entire contraction, while ultrasound images were detected for eight seconds in the middle of the acquisition. The ultrasound and sEMG recordings were synchronised based on an external generator (StimTrig; LISiN, Politecnico di Torino, Italy) [24].

The sEMG grid (8×8 electrodes, 10 mm inter-electrode distance) was placed on the muscle belly after appropriate skin preparation [25]. Monopolar EMG signals were sampled at 2048 Hz through a wireless sEMG acquisition system (MEACS, LISiN, Politecnico di Torino, Turin, Italy) [26]. After band-pass filtering (20-400 Hz), the signals were decomposed into individual MU spike trains [27]. Based on the previous study [14], we selected the MU spike train based on two criteria for each sEMG recording. 1) The centroid of the EMG amplitude distribution closest to an ultrasound-decomposed component (in the lateral direction). 2) A spike-triggered averaged twitch should have a high signal-to-noise (non-zero peak) for the same component.

The ultrasound probe (Verasonics L11-5v with a centre frequency of 7.8125 MHz) was placed in a cross-sectional view in the middle of the transparent sEMG grid [28]. The radio frequency data was recorded using Verasonics Vantage 128 (Verasonics, Inc., Kirkland, WA) sampled at 31.25 MHz with a frame rate of 2500 Hz due to single plane wave imaging. Each dataset comprised 20400 frames (2176×128 pixels each with a field of view equal to 53×40 mm).

After delay-and-sum beamforming, the regions after the maximum sEMG range were removed resulting in a 20×40 mm field of view (850×128 pixels) and the dataset was downsampled to a frame rate of 1250 Hz. Each pixel was filtered over time with a 1D median filter with an order equal to 10 ms (12 samples). Displacement velocity images were calculated based on the radio frequency data using 2D autocorrelation velocity tracking [18] with 0.5 mm (20 samples) indepth and a sliding window of 10 ms (12 frames). The regions were downsampled axially by a factor of thirteen, resulting in 10200 frames where each frame was 64×128 pixels (about 0.3×0.3 mm per pixel). Finally, an ROI of 64×64 pixels was selected [19] for further analysis, where the ROI was placed to contain the activity of a MU.

The study was conducted following the Declaration of Helsinki and the procedure approved by the Regional Ethics Committee (Commissione di Vigilanza, Servizio Sanitario Nazionale-Regione Piemonte, Torino, Italy). Informed consent was obtained from all participants after receiving a detailed explanation of the study procedures.

### Rearranging the data matrix from 3D to 2D

The velocity images contain the axial displacement velocities and can be represented as a three-dimensional data matrix ***Y***. By vectorizing the images, the matrix becomes two-dimensional with the size of *n_s_* × *n_t_*, where *n_s_* = *n_x_* × *n_y_*, and *n_x_* and *n_y_* denote the number of pixels for the width and height, and *n_t_* is the number of images.

### Spatiotemporal independent component analysis (stICA)

The stICA decomposition algorithm will be described based on previous papers [7,19]. First, the mean in both dimensions of a two-dimensional data matrix ***Y*** is subtracted. Then, the matrix is decomposed using Singular Value Decomposition (SVD). The first *k* eigenimages and corresponding eigenvectors are selected and decomposed using spatiotemporal independent component analysis (stICA) [29]. We used *k* = 25 for the nEMG recordings [19] and *k* = 50 for the sEMG recordings [14]. The stICA problem can be formulated as ***Y*** = ***S*Λ*T***^⊤^, where *S* is a vectorized spatial matrix (dimension *n_s_* × *k*), ***T*** is a temporal matrix (dimension *n_t_* × *k*), and Λ is a diagonal scaling matrix [29]. ***S*** and ***T*** have been estimated based on a kurtosis-based cost function based on the Hyperbolic secant distribution [7,19]. The algorithm includes a weight (α) to focus on spatial or temporal separation. For example, *α* = 0.5 means that there is equal weight on spatial and temporal cost function in the separation process. Here, we focused on the spatial cost function (*α* = 1.0), which is the highest performance in terms of the rate of agreement with the MU spike trains [11].

### The velocity-based blind source separation algorithm (velBSS)

The proposed decomposition algorithm has a similar pipeline as the one above. First, the mean is subtracted in both dimensions of a two-dimensional data matrix ***Y***. Second, random SVD [30,31] is applied to decompose ***Y*** into *k* eigenimages and corresponding eigenvectors. Random SVD requires two parameters (except parameter *k*, which was selected as described above): an exponent parameter *q* and an oversampling parameter *p*. Here, we chose *q* = 1 and *p* = 5 [30]. Third, ***Y*** is whitened [32] based on the eigenimages and eigenvectors. Finally, a fixed-point iteration scheme is used to decompose ***Y***. The fixed-point iteration can be formulated as ***Y*** = ***ST***^⊤^, where ***S*** is a vectorized spatial matrix (dimension *n_s_* × *k*), and ***T*** is a temporal matrix (dimension *n_t_* × *k*). To estimate ***S*** and ***T***, we have used a skewness-based gradient function with the first derivative equal to *g*(*x*) = *x*^2^ (and *g′* equals the second derivative) [32]. However, a kurtotic distribution as in stICA above should provide similar results. The proposed algorithm only considers spatial separation, which is a good feature for this data type [11]. The overall algorithm is summarized in pseudo-code below:

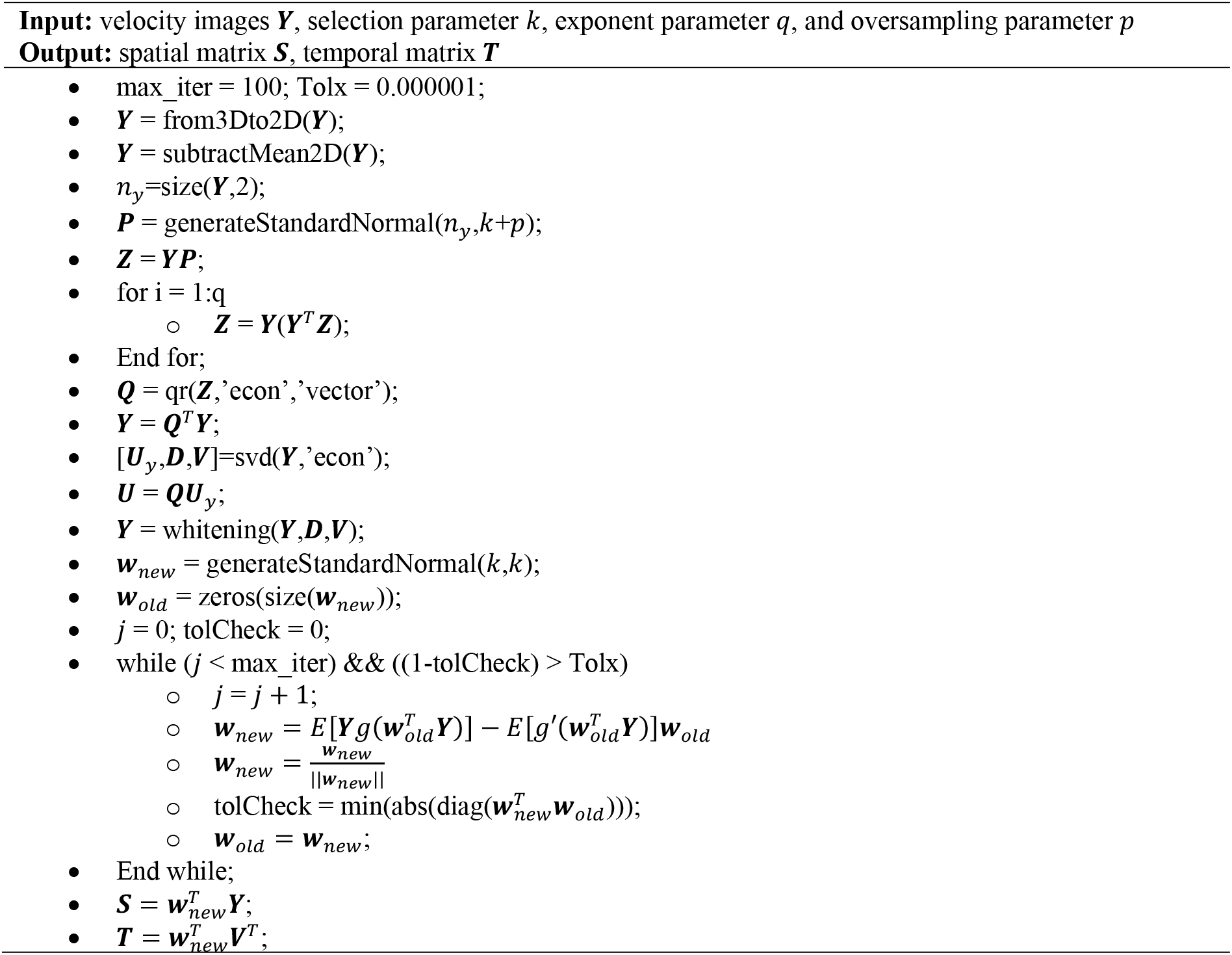

### Comparison between algorithm outputs

The output from the algorithms will be *k* spatial maps and *k* temporal responses (spatiotemporal components), where *k* = 25 for nEMG recordings and *k* = 50 sEMG recordings (see above).

We already know which components are associated with MU activity from nEMG and sEMG using stICA because the raw data originated from the two previous studies [14,19]. Therefore, the components (from velBSS) were matched with the ones from stICA based on finding the components with the highest spatial correlation (vectorized images) and temporal correlation. Then, these matched components were assessed by correlation and presented for visual assessment.

The computational time (and all other data processing and analysis) was performed using MATLAB (2022a, MathWorks, Natick, MA, USA) on an iMac 4.2 GHz Intel Core i7 with 64 GB 2400 MHz DDR4. Ten independent iterations were run, and their mean value was presented.

## Results

### Spatiotemporal consistencies

The (vectorized) spatial maps (***S***) and temporal responses (***T***) extracted from stICA and velBSS were highly correlated for the six paired outputs (0.86 ± 0.05 and 0.87 ± 0.06). In addition, the six paired spike-triggered averaged twitch responses (using the MU spike trains from nEMG and sEMG) had an extremely high correlation (0.99 ± 0.01) based on 19.3 ± 4.0 firings (nEMG) and 90.0 ± 10.6 firings (sEMG), respectively. The spatial maps and the triggered twitch responses for all recordings are shown in Fig. 2.

**Figure 2.**
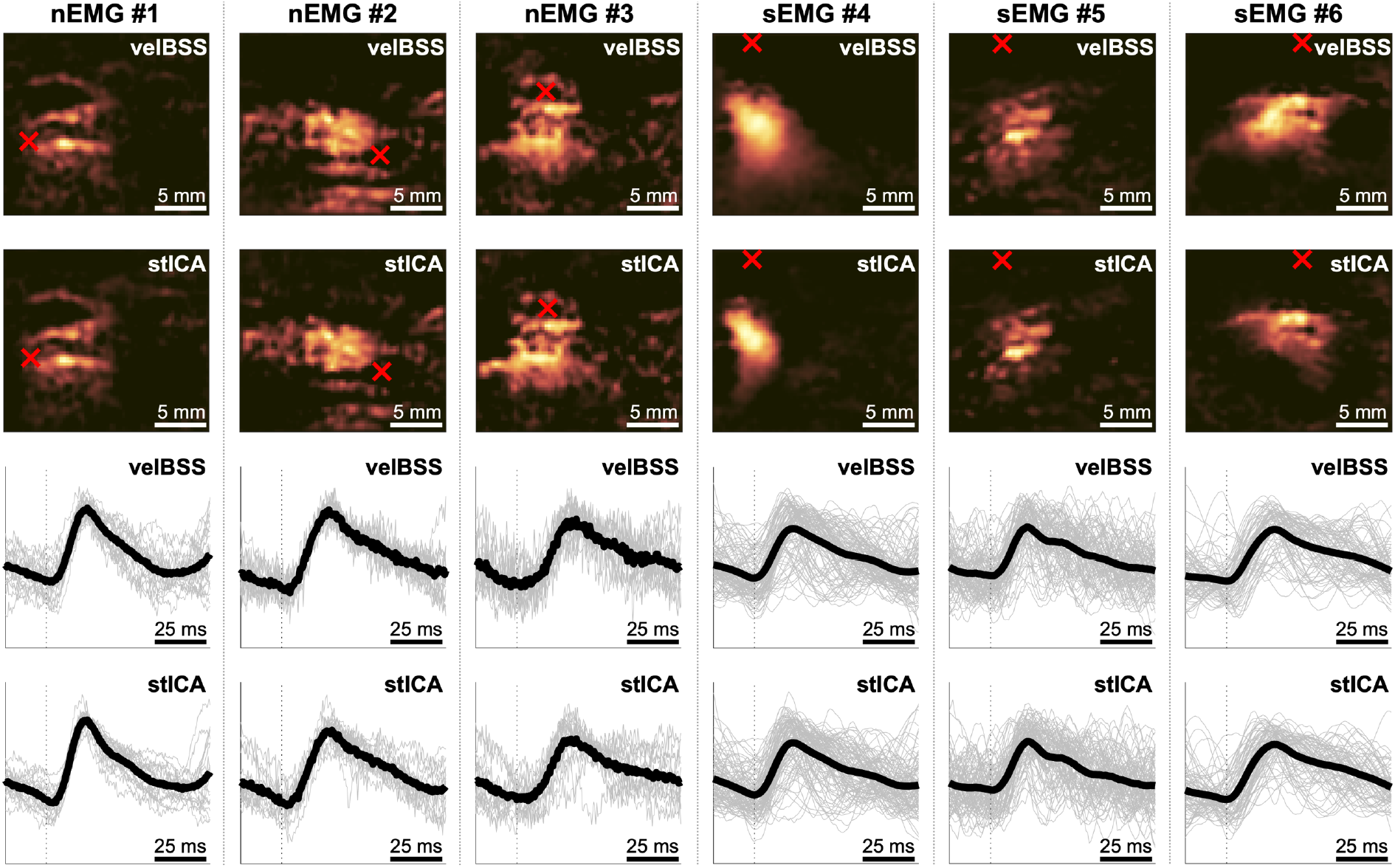
The six selected spatial maps (first two rows) and the corresponding spike-triggered averaged twitch responses (last two rows) from displacement velocity images associated with single motor units (MUs) from needle electromyography (nEMG) and surface EMG (sEMG). The needle tip location (nEMG) and the centroid of the EMG amplitude distribution (sEMG) are illustrated with red crosses. The components were extracted using 1) spatiotemporal independent component analysis (stICA) and 2) velocity-based blind source separation (velBSS). The decomposed results were highly correlated for the six paired and vectorized outputs (0.86 ± 0.05 and 0.99 ± 0.01). Note that the spike-triggered averaged twitch responses denote the average temporal responses of decomposed components triggered using MU spike trains (black vertical dotted line) from nEMG (19.3 ± 4.0 firings) or sEMG (90.0 ± 10.6 firings).

### Computational time

The proposed method (velBSS) had a significantly lower computational time than stICA (Table 1). velBSS decomposed the matrix from the nEMG reference recordings (size 64×64×4012) into 25 components in about 0.17 s compared to stICA, which needed about 18-19 seconds. For the sEMG reference recordings, velBSS decomposed the matrix (size 64×64×10189) into 50 components in about 0.45 s compared to stICA, which needed about 26-30 seconds. Given this, velBSS was at least 50 times faster than stICA.

**Table 1.** The computational time to decompose the data matrix for different recordings and subjects.

|  | Method 1: stICA | Method 2: velBSS |
| --- | --- | --- |
| **nEMG recordings\*** |  |  |
| Subject 1 | 19.15 seconds | 0.17 seconds |
| Subject 2 | 18.92 seconds | 0.17 seconds |
| Subject 3 | 18.72 seconds | 0.17 seconds |
| **sEMG recordings\*\*** |  |  |
| Subject 4 | 26.79 seconds | 0.45 seconds |
| Subject 5 | 30.50 seconds | 0.44 seconds |
| Subject 6 | 26.37 seconds | 0.45 seconds |

stICA = spatiotemporal independent component analysis. velBSS = velocity blind source separation. nEMG = needle electromyography. sEMG = surface electromyography. \* The data dimension was 64x64x4012 and decomposed into 25 components. \*\* The data dimension was 64x64x10189 and decomposed into 50 components. The computational time is based on ten iterations.

## Discussion

This study proposed a fast BSS algorithm (velBSS) suitable for online purposes that decomposes velocity images from low-force voluntary isometric contractions into putative MU activities. The proposed algorithm was compared against stICA in six different recordings, including two ultrasound research systems, six subjects, and two EMG systems (needle vs surface electrodes). We found that the spatial and temporal correlation between the MU-associated components from velBSS and stICA was high and that the computational time for velBSS was at least 50 times less than for stICA.

The proposed velBSS algorithm requires three parameters: the number of components *k* to be decomposed based on the data matrix and the two random SVD parameters *q* and *p* associated with the exponent and oversampling. Although these parameters could have been optimised further, a change in the parameters would only occur for higher correlation values.

The computational time for velBSS was about half a second for an eight-second-long dataset at 1250 Hz. This time could be optimised further using parallelisation and a window-based procedure of smaller time windows. For example, using the Hungarian method [33], one may use the estimated projection vector from the former window to initialise the current window and map the two projection vectors. Nevertheless, the algorithm takes velocity images as input. The steps from plane wave transmission to processed velocity images remain to be solved, where GPU processing will play an important role in minimising the beamforming and estimation of velocity images.

In conclusion, the present algorithm (velBSS) outperforms the currently available method (stICA). It provides a promising translation towards an online pipeline and will be important in the continued development of this research field of functional neuromuscular imaging.

## Acknowledgements

Thanks to Alberto Botter and Marco Carbonaro at Politecnico di Torino, Italy, for allowing us to use a subset of their data in this study. This research received no specific grant from public, commercial, or not-for-profit funding agencies.

## Notes

### Competing Interest Statement

The authors have declared no competing interest.

